# A live-imaging system for Arabidopsis leaf primordia at early stages

**DOI:** 10.64898/2026.02.08.704715

**Authors:** Yujie Zhao, Hokuto Nakayama, Satohiro Okuda, Tetsuya Higashiyama, Hirokazu Tsukaya

## Abstract

Live imaging is one of the most powerful methods to reveal the morphogenesis of plant organs. However, the highly three-dimensional structure of plant organs always poses technical challenges. For example, the basal region of leaf primordia is rarely observed because of the shape of leaf primordia and the sudden shift in geometry at the point where the leaf primordium connects to the hypocotyl. In this work, we developed a new live-imaging system that is suitable for observing the developmental process of the basal region of Arabidopsis leaf primordia at early stages. Using this system, we achieved continuous observation of the basal region of early Arabidopsis leaf primordia for more than 50 hours.

## Introduction

Live imaging is a key approach for uncovering the detailed morphogenesis of plant organs. With advances in live imaging systems, dynamic cellular processes can now be visualized in an increasing variety of plant tissues (Calder et al., 2015; Mizuta et al., 2024; Yadav & Roeder, 2024). For instance, the live-imaging system described by Yadav and Roeder (2024) enabled detailed observation of cellular dynamics within sepals in *Arabidopsis thaliana* (hereafter, Arabidopsis). However, the current systems are often constrained by the complex three-dimensional morphology of target organs. For example, leaf primordia exhibit a crescent-shaped morphology in the medio-lateral plane (Watanabe & Okada, 2003), and the hypocotyl is wider than the leaf primordium in the radial direction (Fig. 1). As a result, the basal region of the leaf primordium is positioned within the recessed area between the stem and the primordium itself. This configuration requires greater imaging depth and poses significant challenges for effective observation. Although multiple approaches have been reported to observe the curved surface of plant tissues (Silveira et al., 2021; Harline & Roeder, 2023), these methods still have several limitations. Harline & Roeder (2023) described a system that flattens the curved surface by directly suspending a coverslip above the sample. This system is primarily limited to observing the middle part of the abaxial side of the leaf primordium and has difficulty accessing the region hidden within the recessed area between the stem and the primordium. In addition, many of these systems require a confocal microscope with deep imaging depth.

**Figure 1.**
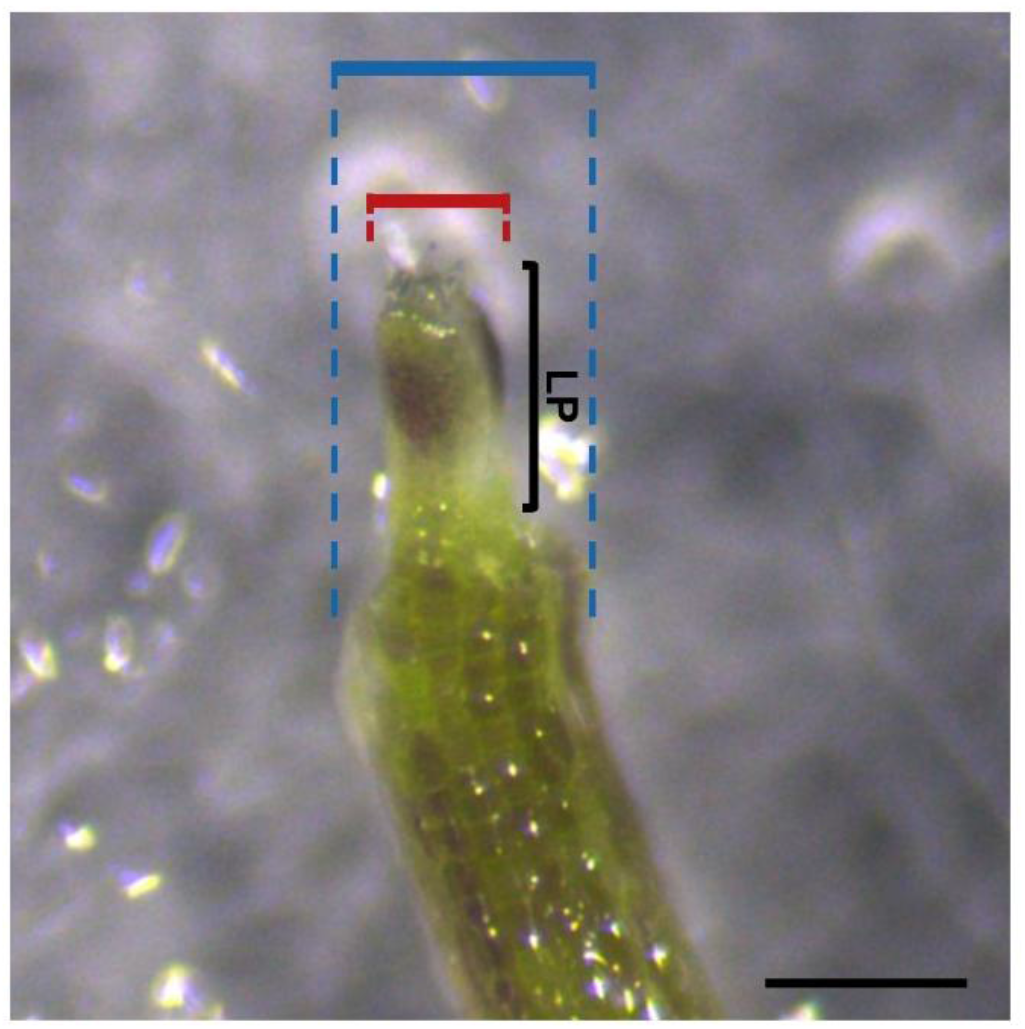
Structure of the junction of leaf primordium and hypocotyl. The image shows the tip part of a 3.5 DAS seedling. The red and blue lines show the width of leaf primordia and hypocotyl, respectively. LP: Leaf Primordium. Scale bar = 200 μm.

In the current study, we developed a new live-imaging system that extends previously published approaches to enable imaging of the basal region of early Arabidopsis leaf primordia. Additionally, this system does not require a microscope with deep imaging depth. We believe that this system has the potential to be applied to a wide range of plant tissues to provide detailed growth data.

## Materials and methods

### Plant material and growth condition (see Table 1 for materials and consumables)

The *ML1*_*pro*_*:mCitrine-RCI2a ML1*_*pro*_*:H2B-TFP* (Robinson et al., 2018; ABRC accession CS73343) was used in this work. Seeds were sterilized in 1% (v/v) sodium hypochlorite solution containing 0.1% (v/v) Triton X-100 on a mini rotator (ARC-100, AS ONE Corporation, Osaka, Japan) for 2-3 mins. Sterilized seeds were washed in autoclaved MilliQ water 2-3 times and sown on 1/2 Murashige and Skoog media (Murashige & Skoog, 1962), supplemented with 1% (w/v) agar, 1% (w/v) sucrose, and adjusted to pH 5.7. After stratification at 4°C for 2-3 days, the seedlings were grown in a closed growth chamber set at 22°C under approximately 50 μmol m^−2^ s^−1^ of continuous light from LED lamps (HotaluX, Tokyo, Japan).

**Table 1.** Materials and consumables used in this study.

| Table 1. Materials and consumables used in this study |  |
| --- | --- |
| Materials / Consumables | Source |
| Murashige and Skoog plant salt mixture | SHIOTANI M.S. CO., LTD., Amagasaki, Japan |
| Agar | FUJIFILM, Osaka, Japan |
| Sucrose | NACALAI TESQUE INC., Kyoto, Japan |
| Triton-X100 | NACALAI TESQUE INC., Kyoto, Japan |
| Sodium hypochlorite solution | NACALAI TESQUE INC., Kyoto, Japan |
| 26 GX 1/2 needle | TERUMO, Tokyo, Japan |
| 35 mm/glass base dish (glass 27 Ø) | AGC TECHNO GLASS CO., LTD., Shizuoka, Japan |
| Coverslip | MATSUNAMI GLASS IND., LTD., Osaka, Japan |
| Silicone film | AS ONE Corporation, Osaka, Japan |
| Vacuum grease | DDP Specialty Electronic Materials US 9, LLC, Hemlock, USA |
| Perfluorodecalin | Sigma-Aldrich, St. Louis, USA |
| Hyponex | HYPONeX, Osaka, Japan |
| Rockwool | Grodan, Roermond, Netherlands |

To observe whether seedlings grow normally, several seedlings were moved to rockwool after live imaging and watered with an adequate amount of Hyponex solution (1 g/L) every 2-3 days. All other growth conditions were as described above.

### Imaging condition

The seedlings were imaged by using an inverted fluorescence microscope (IX81DC2; Olympus Corporation, Tokyo, Japan) equipped with a spinning disk-scan confocal system (CSU-X1; Yokogawa Electric, Tokyo, Japan), a 488-nm LD laser (Sapphire; Coherent, Santa Clara, USA), and an EM-CCD camera (Evolve 512; Photometrics, Tucson, Arizona, USA). A 40 × water immersion objective lens (UApo/340 40×/1.15 Water, WD = 0.25 mm, NA = 1.15; Olympus) was used. The fluorescent signal was excited at 488 nm and was detected with a band-pass filter, 520-535 nm. Samples were observed every two hours. After imaging, the samples were returned to the growth chamber (22°C, with approximately 50 μmol m^−2^ s^−1^ of continuous light). For each observation, the focus and z-range were manually adjusted.

## Results

### Workflow of live imaging

Previous live-imaging approaches are limited in their ability to access complex 3D-shaped structures, such as the basal region of leaf primordia, and highly rely on the capability of microscopes with deep imaging depth. To tackle this limitation, we developed a new live-imaging system that enables stable and long-term live imaging observation with regular confocal microscopes. At this time, we used the basal region of leaf primordia as a sample that is difficult to image. Our system exposes tissue located within the recessed region by using vacuum grease to closely attach the sample to the bottom of the culture dish. In addition, the observation region is immersed in perfluorodecalin (PFD; Table 1), while culture medium is supplied only to the root region. PFD was selected for its ability to provide high image quality and was likely to help maintain the growth of seedlings due to its high oxygen-dissolving capacity (Littlejohn et al., 2010). Together, these features reduce the requirement for deep imaging capability, enabling observation with regular microscopes. The overall workflow of the system is outlined below (Fig. 2, see Table 1 for materials and consumables).

**Figure 2.**
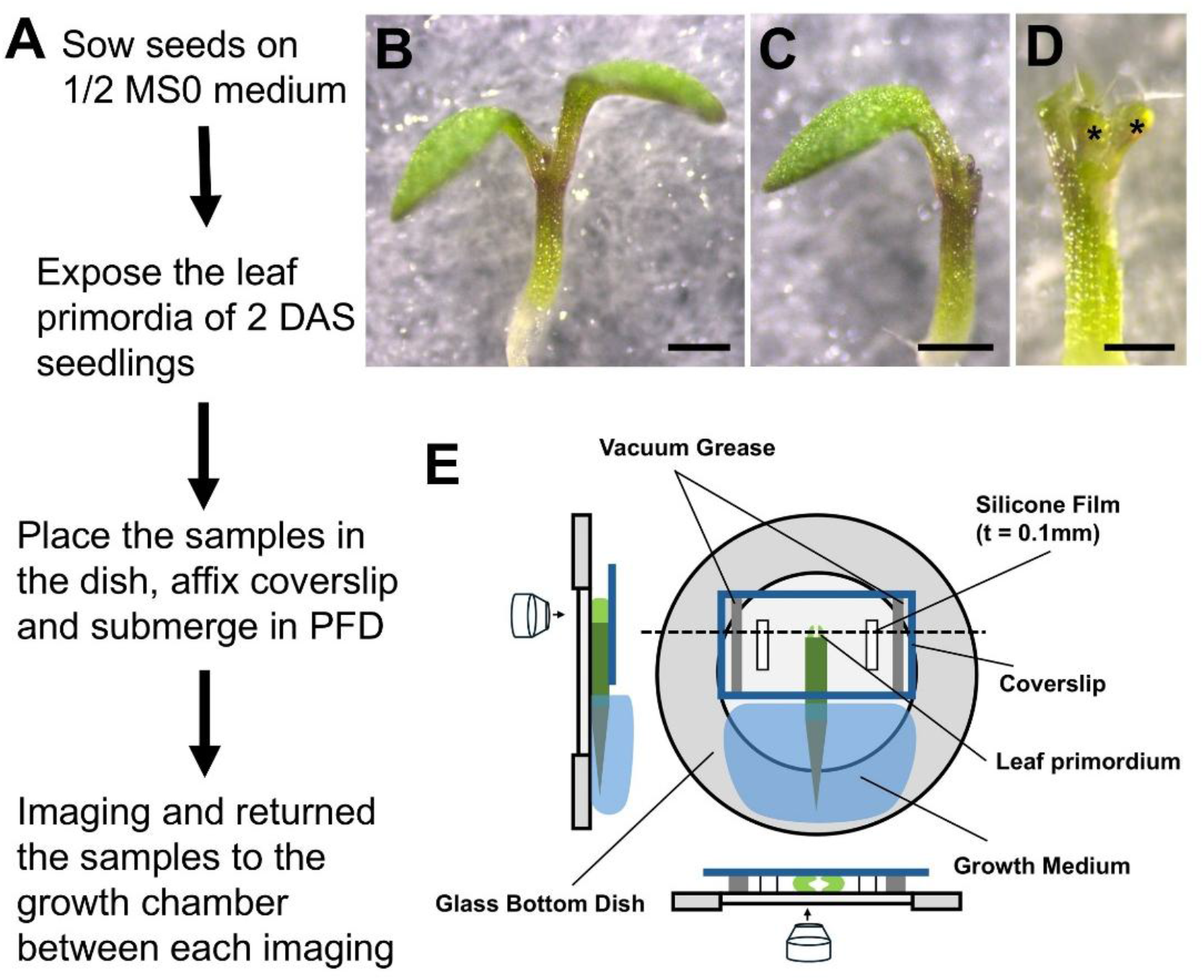
An imaging system developed in this study. (A) The workflow for the configuration of the newly developed live-imaging system. (B-D) The process of exposing leaf primordia. (B) One 3.5 DAS seedling prior to cotyledon excision (C) One cotyledon is removed by using the needle and small brush. (D) The other cotyledon was removed in the same manner as described above, except that the petiole. The seedling is prepared for live imaging observation. The black asterisks highlight two leaf primordia. Scale bars = 500 μm in (B and C), 250 μm in (D). (E) Images of sample mounting in the newly developed live-imaging system. The dashed line represents the plane shown in the front view as projected onto the top view. The remaining basal part of cotyledons after dissection is not depicted in the model to highlight the leaf primordia.

1. 2-2.5 days after stratification (DAS), seedlings of *ML1*_*pro*_*:mCitrine-RCI2a ML1*_*pro*_*:H2B-TFP* were used for live imaging. In this line, the plasma membrane and nuclei are labeled, respectively, enabling visualization of epidermal cells without staining.
2. One cotyledon was dissected off at the base of the seedling by a needle (Table 1) to expose the leaf primordia for observation (Fig. 2B and C).
3. Another cotyledon was dissected off in the same way, except for the petiole part, which can facilitate orientation adjustment of the seedlings and get an appropriate image (Fig. 2D).
4. Dissected samples were placed in glass base dishes (Table 1) and affixed with coverslips, which were adjusted to a suitable size. Between the bottom of glass base dishes and the coverslips, two strips of silicone film (thickness = 1 mm; Table 1) were placed to provide sufficient space for the seedlings and prevent tissue damage, and an adequate amount of vacuum grease (Table 1) was placed to ensure the plant remained closely attached to the bottom of glass base dish, improving imaging quality. These setups allowed us to transform the curved surface of the leaf primordia into a horizontal plane, enabling detailed observation with more cells during the developmental process (Fig. 2E).
5. The remaining space between the bottom of glass base dishes and coverslips was filled with PFD.
6. Some 1/2 Murashige and Skoog media, which were supplemented with 1% (w/v) agar, 1% (w/v) sucrose, and adjusted to pH=5.7, were placed on the seedlings’ roots to provide water and nutrients (Fig. 2E).
7. Observe samples with confocal microscope. During the observation intervals, the samples were returned to the growth chamber to allow continued growth.

### Imaging of the basal region of early Arabidopsis leaf primordia

By using our live-imaging system, we could observe the developmental process of multiple leaf primordia for more than 50 hours from the same imaging perspective, one representative example is shown in Fig. 3. During this experiment, the length of leaf primordia increased, and cell expansion and active cell proliferation were observed well (Figs. 3 and 4), suggesting that leaf primordia grow under the newly established system. In addition, samples used for live imaging were able to resume normal growth after the experiment (Fig. 5).

**Figure 3.**
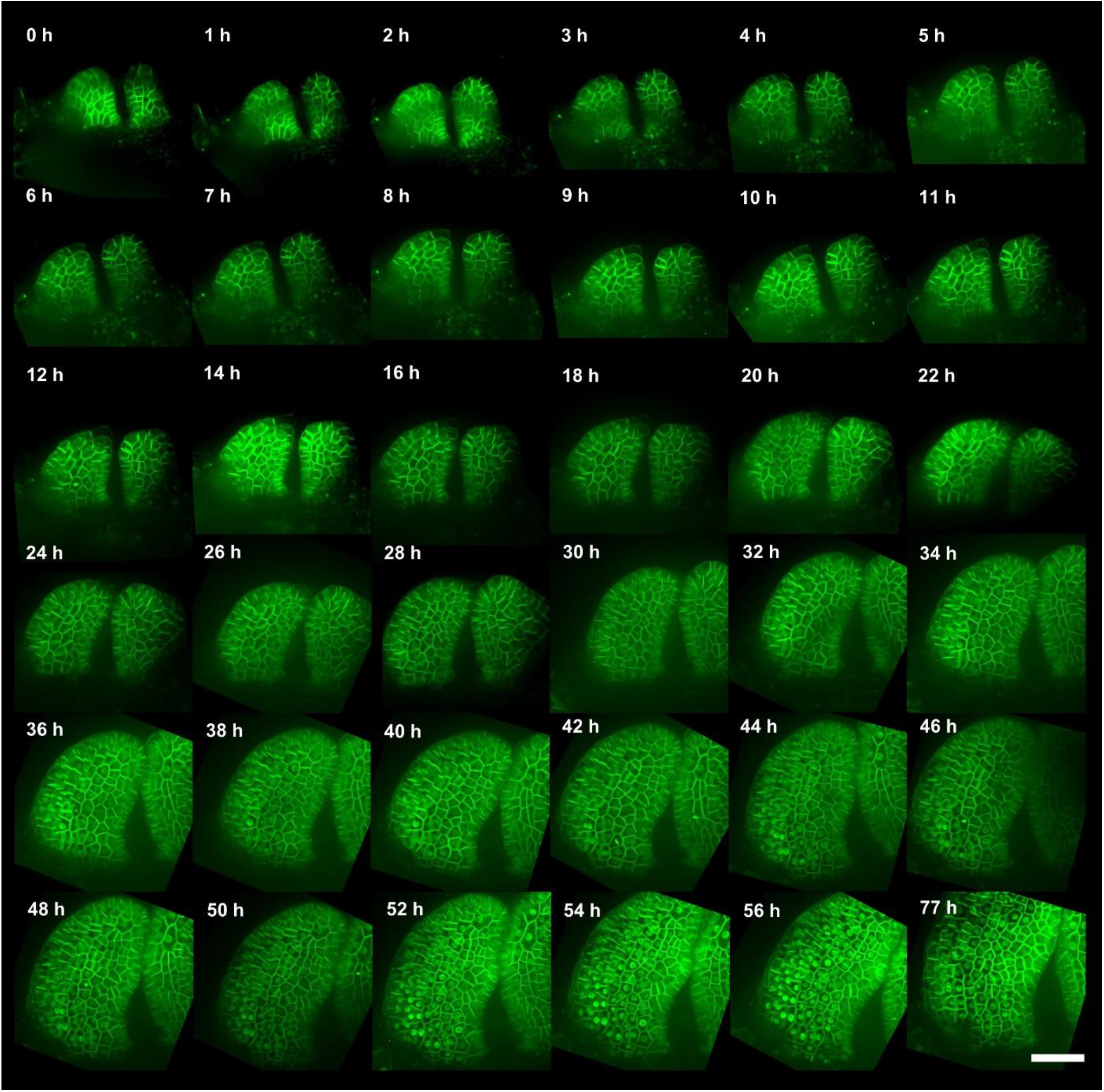
Live imaging data with leaf primordia. The ImageJ z-projected images of the identical leaf primordia of *ML1*_*pro*_*:mCitrine-RCI2a ML1*_*pro*_*:H2B-TFP*, which labeled the plasma membrane and nucleus of the outermost epidermal cells in meristem tissue, respectively, captured from 2.5 to 5.5 DAS. Images corresponding to 0 h to 77 h after the start of semi-live imaging are shown sequentially. Scale bar = 50 μm.

**Figure 4.**
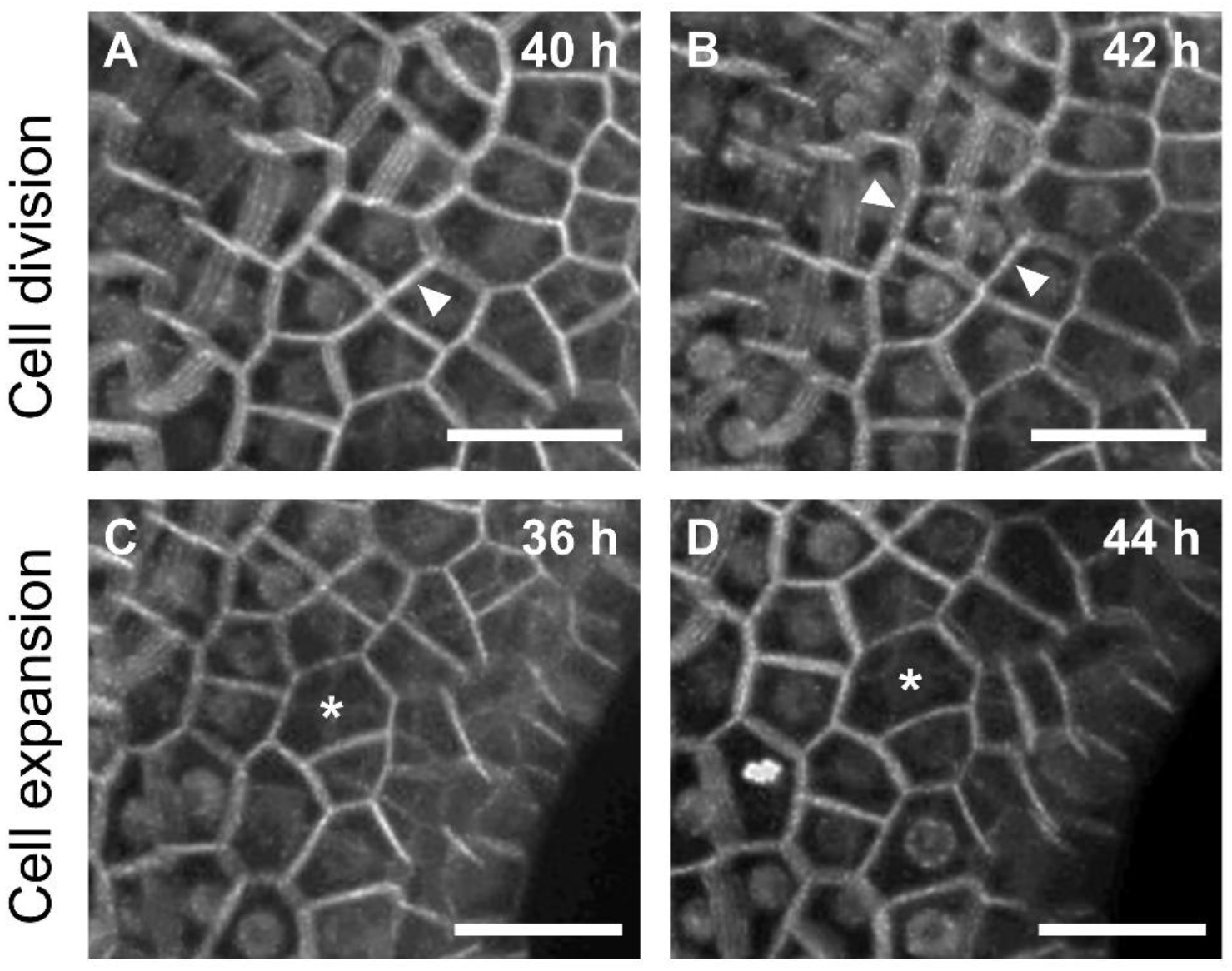
Examples of cell division and cell expansion during live imaging. The images show the cell division observed between 40 h and 42 h (A and B), and cell expansion observed between 36 h and 44 h (C and D) after the start of live imaging. The leaf primordium of *ML1*_*pro*_*:mCitrine-RCI2a ML1*_*pro*_*:H2B-TFP* was observed, which labeled the plasma membrane and nucleus of the outermost epidermal cells in meristem tissue, respectively. The white arrowheads indicate the cells that are undergoing division. The asterisks highlight the cells that are undergoing expansion. Scale bars = 20 μm.

**Figure 5.**
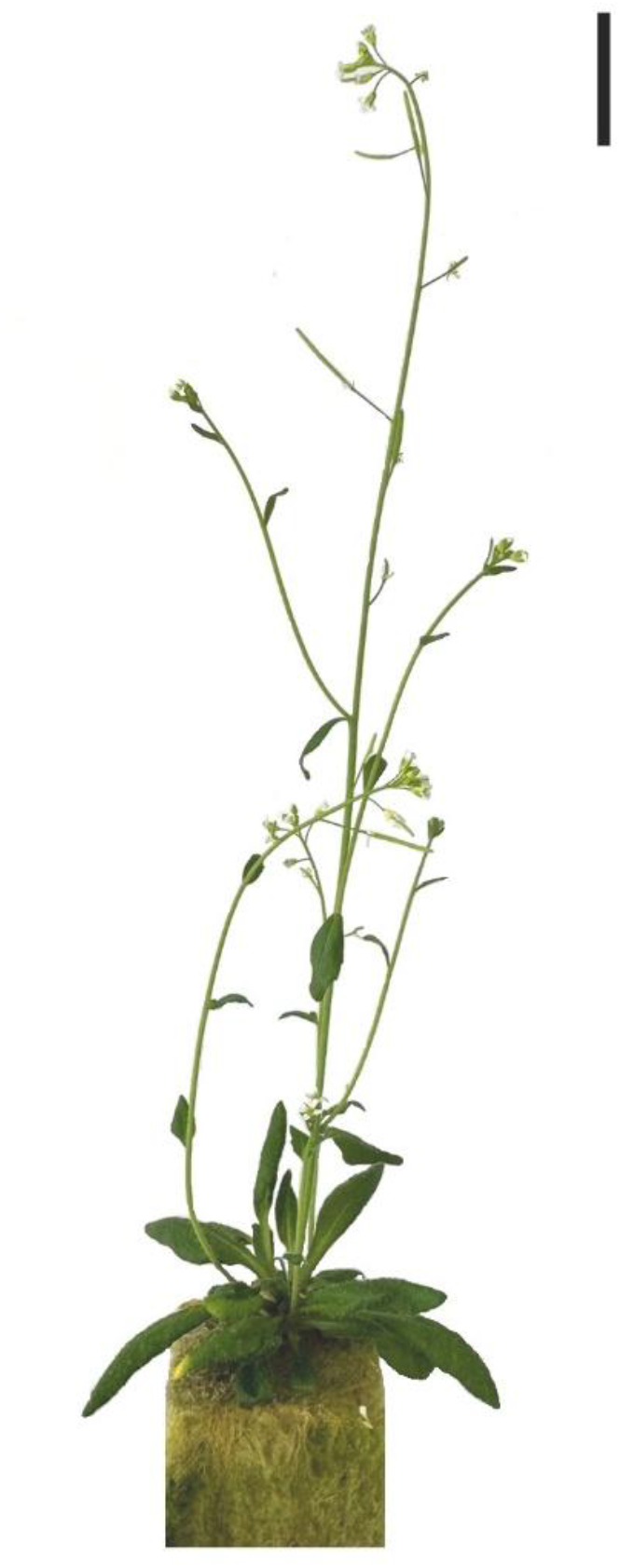
Example of post-imaging growth of Arabidopsis seedling. The image shows the seedling three weeks after live imaging. Scale bar = 2 cm.

## Discussion

In this work, we developed a live-imaging system that allows long-term observation of complex three-dimensional structures, including the basal region of Arabidopsis leaf primordia at early stages, which has been difficult to access with previous methods. Notably, our live-imaging system is simple and does not require expensive hardware modules or rely on complex experimental procedures. Moreover, our results demonstrate that the system supports sustained growth of leaf primordia during long-term imaging and enables continuous observation of early petiole development over extended periods using a conventional inverted microscope, such as the Olympus IX81. By combining a simple mounting strategy with intermittent imaging, the system reduces the requirement for imaging depth and provides a stable and reproducible imaging perspective over time. This approach could be extended to other plant tissues, although additional experiments are necessary to confirm its broader applicability.

## Acknowledgements

This work was supported by JSPS KAKENHI (JP19K23742, JP20K06682, JP20KK0340 to HN) and a Grant-in-Aid for Scientific Research on Innovation Areas (JP19H05672 to HT).

## Author Contributions

Y.Z.: System design, Validation, Writing–original draft, Writing – review & editing. H.N.: System design, Writing – review & editing, Supervision, Funding acquisition. S.O.: System design, Resources, Writing – review & editing. T.H.: Resources, Writing – review & editing. H.T.: Funding acquisition, Supervision, Writing – review & editing.

## Notes

### Competing Interest Statement

The authors have declared no competing interest.

